# The effect of psychedelics on associative learning: a systematic review

**DOI:** 10.1101/2025.07.21.665870

**Authors:** Alice Caulfield, Lorraine Li, Famia Askari, Clara Belessiotis-Richards, Allan H Young, Mitul Mehta

**Author notes:** **CORRESPONDING AUTHOR:** Alice Caulfield. Centre for Neuroimaging Sciences, De Crespigny Park, Camberwell, SE5 8AF.

## Abstract

**Introduction:** Psychedelics are emerging as potential treatments for neuropsychiatric conditions, with evidence suggesting a single administration can lead to enduring behavioural changes. While the underlying putative mechanism(s) remain unclear, there is evidence supporting altered learning as a key candidate.

**Aim:** This systematic review examined studies assessing the effects of psychedelics on associative learning in both humans and animals.

**Methods:** Electronic databases were searched up until 13/01/2025 for studies investigating any difference in learning after psychedelic administration.

**Results:** 31 studies were included (29 in animals, 2 in humans). Classical and operant conditioning paradigms were employed, including fear extinction, conditioned avoidance, and reversal learning. Studies assessed acute and post-acute effects, however repeated dosing paradigms often obscured this distinction. There was considerable heterogeneity in study designs, paradigms, drug administration timings and doses, and behavioural effects appeared to be influenced by dose, timing, training intensity, and sex. Due to between-study heterogeneity, a meta-analysis was not possible.

Evidence suggests that psychedelic administration enhances associative learning in animals across paradigms, although findings were not entirely consistent. Possible mechanisms identified were increased prediction error sensitivity, serotonin receptor agonism, and structural plasticity. Learning enhancements may extend into the post-acute phase and appear to depend on active environmental engagement during this window.

**Conclusion:** Studies suggest that psychedelics enhance associative learning in animals; however, these findings are yet to be translated into humans. Understanding whether a period of enhanced learning follows the psychedelic experience may have important implications for psychedelic-assisted psychotherapy, where behavioural changes must generalise and persist beyond the drug-induced state.

## INTRODUCTION

Psychedelics are a class of psychoactive substances which induce altered states of consciousness by modulating serotonergic systems, primarily through 5-HT_2A_ receptor agonism, with profound effects on perception, cognition, and mood (Madsen et al., 2019; Preller et al., 2018; Doss et al., 2021; Pokorny et al., 2020). Research into psychedelics was widely prohibited in the mid-20th century, following their association with countercultural movements (Dyck, 2008), however since the early 2000s, there has been a resurgence of scientific interest, with a corresponding growing body of evidence suggesting that psychedelics may be effective treatments for a range of mental health conditions (Andersen et al., 2021). A phase 3 trial investigating psilocybin vs placebo as a treatment for treatment-resistant depression has recently shown a small but significant improvement in symptoms (ClinicalTrials.gov, 2025).

The observed improvements in clinical studies (Griffiths et al., 2016; Palhano-Fontes et al., 2019; Carhart-Harris et al., 2018a; Sanches et al., 2016; Moreno et al., 2006; Ross et al., 2016; Gasser et al., 2014; Grob et al., 2011; Bogenschutz et al., 2015; Johnson et al., 2017; Davis et al., 2021), and sustained changes following psychedelic administration (Griffiths et al., 2018; Griffiths et al., 2011; Griffiths et al., 2006; Doss et al., 2021; Lebedev et al., 2016; Schmid and Liechti, 2018) suggest that a single dose of a psychedelic may result in long-lasting changes, at either a behavioural or neural level. However, we lack an understanding of why this occurs and consequently, how putative therapeutic effects may be augmented. A candidate mechanism of importance is psychedelic-induced enhanced neuroplasticity (for reviews, see (Calder and Hasler, 2023; De Vos et al., 2021; Olson, 2022), however multiple putative mechanisms of psychedelic action have been identified and may operate synergistically. At a receptor-level, classical serotonergic psychedelics are 5-HT_2A_ receptor agonists, however the receptor binding profiles are far more complex (for review, see (Halberstadt and Geyer, 2011)). Many other processes have been identified that underlie psychedelic-induced neuroplasticity cascades (Ly et al., 2018), including TrkB receptor activation (Moliner et al., 2023), molecular-level changes such as increased expression of c-Fos (Nichols and Sanders-Bush, 2002; Frankel and Cunningham, 2002), altered gene expression (Nichols and Sanders-Bush, 2002) and remodelling of the extracellular matrix (Nardou et al., 2023). Systems-level changes within a computational framework include a relaxation of prior beliefs in the context of predictive coding, which has dominated the recent literature (Carhart-Harris and Friston, 2019). Behaviourally, changes in cognitive flexibility (Doss et al., 2021; Wießner et al., 2022), creativity (Prochazkova et al., 2018), and enduring changes in wellbeing and personality (Aday et al., 2020) have been described. How these levels may interact, or operate synergistically, remains unclear.

The acute psychedelic experience is highly influenced by the context, both the external environment (setting), and the internal psychological context (the set) (Carhart-Harris et al., 2018b). The quality of the acute experience can affect whether sustained effects are positive (Roseman et al., 2018) or negative (Carbonaro et al., 2016; Halpern et al., 2018). In experimental animals, psychedelics have shown neuroplastic effects (Ly et al., 2018; Nardou et al., 2023), and it is possible that these facilitate sustained behavioural change. However, the relationship between neuroplasticity and behavioural change remains unclear and is challenging to measure, especially in humans. This interaction may be explored by examining learning, which is the process of changing behaviour following the acquisition of new knowledge or experience. There is a long history of evidence for changes in associative learning (see Boxes 1 and 2) under psychedelics; however, this has been largely unexplored in modern-era psychedelic research. The aim of this systematic review is to synthesise studies relating to the effect of psychedelic administration on learning, to understand whether this could be a putative mechanism (among others) of psychedelic action.

## METHODS

### Box 1: Associative learning – a background

Associative learning is the cognitive process by which an individual forms associations between two stimuli, or between an action and its consequence, based on their co-occurrence (for review, see (Pearce and Bouton, 2001)). In practical, real-world contexts, associative learning simply involves linking one factor with another through an individual’s lived experiences, and is a key mechanism through which complex behaviour arises, with important implications for social development and future mental health (Sheridan et al., 2018; Reeb-Sutherland et al., 2012).

Since the work of Pavlov and Thorndike (Thorndike, 1898; Pavlov, 2010), the field has expanded to incorporate mathematical frameworks such as the Rescorla-Wagner (RW) model (Rescorla, 1972) to explain more complex learning processes. The RW model proposes that the strength of associative learning depends on the discrepancy between predicted and actual outcomes, known as prediction error or ‘surprise.’ As associations become stronger, surprise diminishes, correspondingly reducing the rate of learning. This has paved the way for more sophisticated frameworks such as reinforcement learning (RL) and active inference (AI) which are able to incorporate measures of uncertainty allowing for more flexible and adaptive learning in complex, dynamic environments. These advanced models not only account for prediction errors and reward maximization but also enable agents to build and update probabilistic models of their world, optimizing behaviour through continuous interaction and belief updating. This progression is reflected in the literature. Whereas earlier studies used simple task designs with fixed probabilities, newer studies have incorporated probabilistic task designs with computational modelling to characterise richer alterations in learning mechanisms in response to psychopharmacological manipulations.

The various types of associative learning are summarised in box 2, and may be broadly categorised as classical conditioning, or operant conditioning.

### Systematic review protocol

This systematic review was conducted in accordance with the Reporting of Systematic Reviews and Meta-Analyses (PRISMA) guidelines (Page et al., 2021). The search protocol was preregistered on the OSF (ID osf.io/p7yqn). The aim of this systematic review was to compile studies investigating any difference in learning observed after psychedelic administration, relative to a placebo comparison condition or pre-drug condition.

### Search strategy

Three electronic databases were searched (MEDLINE, APA PsychInfo, Embase) for papers up until 13^th^ January 2025, with no restrictions on date. Our search terms were: Learn*.mp AND "psychedelic*" or "hallucinogen*" or "psilocybin" or "4-phosphoryloxy-N,N-dimethyltryptamine" or "psilocin" or "4-hydroxy-N,N-dimethyltryptamine" or "LSD" or "LSD-25" or "lysergic acid diethylamide" or "DMT" or "N,N dimethyltryptamine" or "5-MeO-DMT" or "5-methoxy-N,N dimethyltryptamine" or "ayahuasca" or "2,5-dimethoxy-4-iodoamphetamine". A title restriction was applied to the psychedelic terms only.

### Identification of eligible studies

Studies were included if they compared learning performance after psychedelic exposure vs placebo, in a controlled environment, with a sample size greater than n=6, to reduce the risk of bias due to very small group sizes. Both human and animal studies were included. Review articles, case reports, conference abstracts, and non-English language papers were excluded. After duplicates were removed, all studies returned by the search were screened according to their title and abstract by two independent reviewers (AC and CB-R). Any discrepancies were discussed until resolution. 87 full-text articles were then screened according to the above eligibility criteria.

### Data extraction

Data was extracted following a full-text review of included articles. The following information was extracted: authors, year of publication, study population, psychedelic type, dose, dosing regimen, learning mechanism studied, task paradigm used, and the timing of psychedelic administration relative to tasks, to evaluate whether assessments are during the acute or post-acute period.

### Risk of bias

The risk of bias (ROB) for each study was assessed independently by two authors (LL and FA) using Cochrane’s risk of bias tools: the RoB v2 for RCTs (Sterne et al., 2019) and the SYRCLE tool for animal studies (Hooijmans et al., 2014) and any discrepancies were discussed.

### Synthesis of results

A narrative summary of results is presented, with a tabulated summary of study design. Findings are presented systematically, grouped by type of learning examined (classical conditioning and operant conditioning). All results are presented as compared with a comparable group administered placebo (unless otherwise specified).

## RESULTS

### Study selection

The results of the study selection process are presented in the PRISMA flow chart (Figure 1). 607 records were identified, and after duplicate elimination, 393 records were identified for abstract screening. Of these, 79 articles were found eligible for full-text screening. 45 were excluded after full-text screening, leaving a total of 31 articles included in the final review.

**Figure 1:**
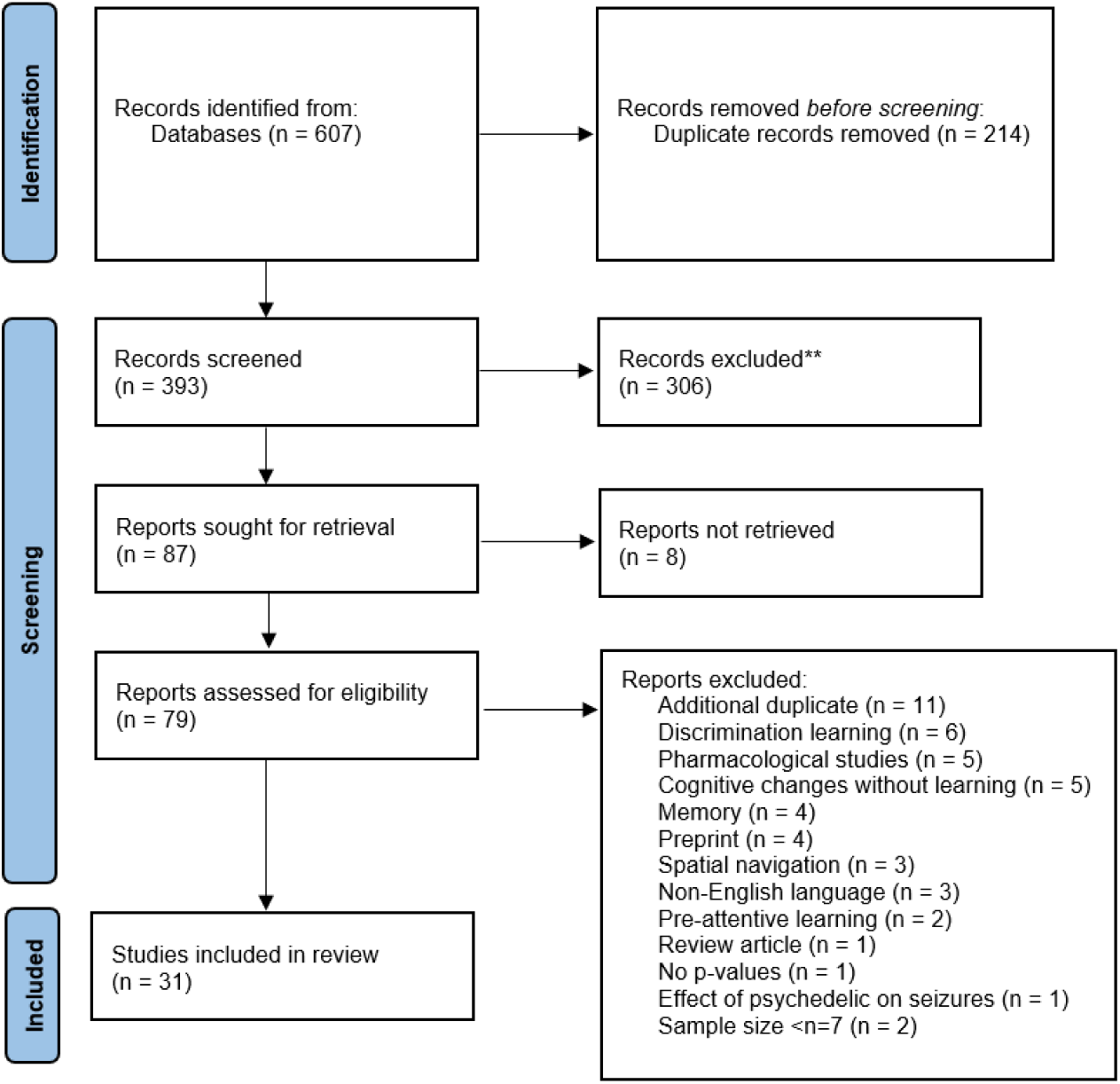
PRISMA flow diagram illustrating the process of selecting studies for the systematic review.

### Study characteristics

Of the 31 studies identified, all but two were conducted in experimental animals, with the remaining two in humans. Psychedelics studied included LSD (n=16), psilocybin (n=8), DOI (n=3), ayahuasca (n=2), 4-OH-DiPT, a psilocybin analogue (n=1), and one study used multiple psychedelics (n=1). Study populations included rats (n=15), mice (n=8), rabbits (n=6) and humans (n=2).

Twelve studies tested learning under the acute drug effect; however, due to daily administration paradigms, interpretation of the relative contributions of acute vs post-acute drug effects was largely obscured after the first session. Twelve studies examined learning during the acute drug effects and were unobscured by possible post-acute effects. Seven studies measured post-acute drug effects and were unobscured by acute effects. All studies assessed associative learning and were categorised based on the paradigm used, as either classical (n=15) or operant conditioning (n=16). For clarity, the summary of results is categorised according to the learning paradigm used: classical acquisition and extinction paradigms; operant paradigms with positive, negative and mixed reinforcement; and reversal learning.

### Risk of bias

The methodological quality of the included studies varied, with inconsistencies across all assessed domains. Of the 29 animal studies evaluated using the SYRCLE tool (Supplementary Table 1), none provided sufficient detail to assess all ten domains. Nine studies (Castellano et al., 1979; Conn et al., 2024; Du et al., 2023; Favaro et al., 2015; Gormezano & Harvey, 1980; Kelly et al., 2024; Pędzich et al., 2022; Romano et al., 2010; Woodburn et al., 2024) were rated as low risk of bias for the domains that were reported, while seven (Butters et al., 1966; Fisher et al., 2024; King et al., 1974; Nardou et al., 2023; Šabanović et al., 2024; Waser et al., 1976; Welsh et al., 1998) had at least one domain rated as high risk. The remaining 13 studies had at least one domain rated as being of some concern, but none were judged to be at high risk of bias.

Two human RCTs were assessed using the RoB 2 tool (Supplementary Table 2). One trial (Casanova et al., 2024) was rated as having a moderate risk of bias due to deviations from the intended intervention, while the other (Kanen et al., 2022) was rated as high risk due to concerns about carryover effects, deviations from the intended intervention, and outcome measurement.

## Results

### Classical conditioning

For a summary of the background to classical conditioning, see Box 2.

#### Box 2: Classical conditioning

##### Acquisition

Classical conditioning acquisition involves associating a neutral stimulus (such as a tone) with an unconditioned stimulus (US) through repeated pairings. The US can be aversive (such as an electrical shock, or an air puff) or appetitive (such as a food reward), and naturally elicits an unconditioned response (UR). Over time, the neutral stimulus becomes a conditioned stimulus (CS), evoking a conditioned response (CR), which resembles the UR, even in the absence of the US. Most studies examining the effect of psychedelics on classical conditioning used an aversive unconditioned stimulus, eliciting a defensive response (see Table 1 for a methodological summary).

##### Extinction

Extinction is the process of learning new associations which compete with the original associations (for reviews see (Bouton, 2004; Myers and Davis, 2002)), and is known to be highly context dependent with limited translation from one context to another (Bouton, 2004). This is observed in fear extinction paradigms with expressions of fear renewal in novel contexts (for review, see (Maren et al., 2013)). The effect of psychedelics on extinction has primarily been studied using fear extinction paradigms.

**Fear extinction** is the learning process whereby exposure to learnt fear-inducing cues, without co-presentation of the aversive stimulus, leads to a reduction in fear response. Fear conditioning paradigms involve co-presentation of a cue, such as a sound, or a particular environment, with an aversive stimulus. This leads to an expression of fear in response to the cue, measured by freezing behaviour. The learnt fear response can be extinguished by repeatedly presenting the conditioned stimulus (CS) without the aversive one, however this is not necessarily driven by forgetting the former association (Gale et al., 2004). Retention can subsequently be tested by measuring freezing behaviour in response to the conditioned stimulus (for review, see (Myers and Davis, 2007). Fear-conditioning is known to display context-dependence (for review, see (Maren et al., 2013)), and therefore, auditory fear conditioning paradigms often test fear extinction in a novel context, to control for the influence of context in initial fear associations.

**Table 1:**
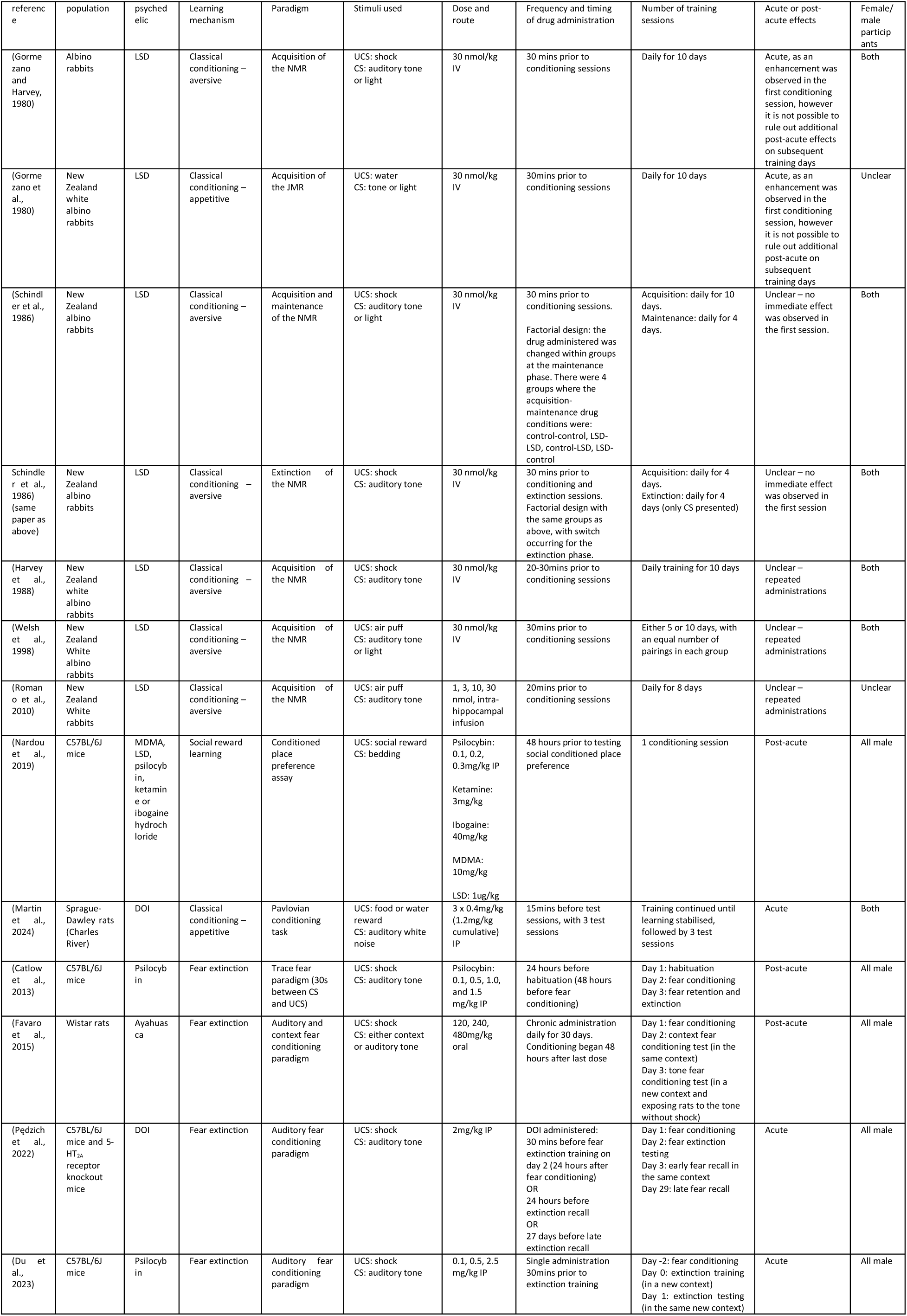
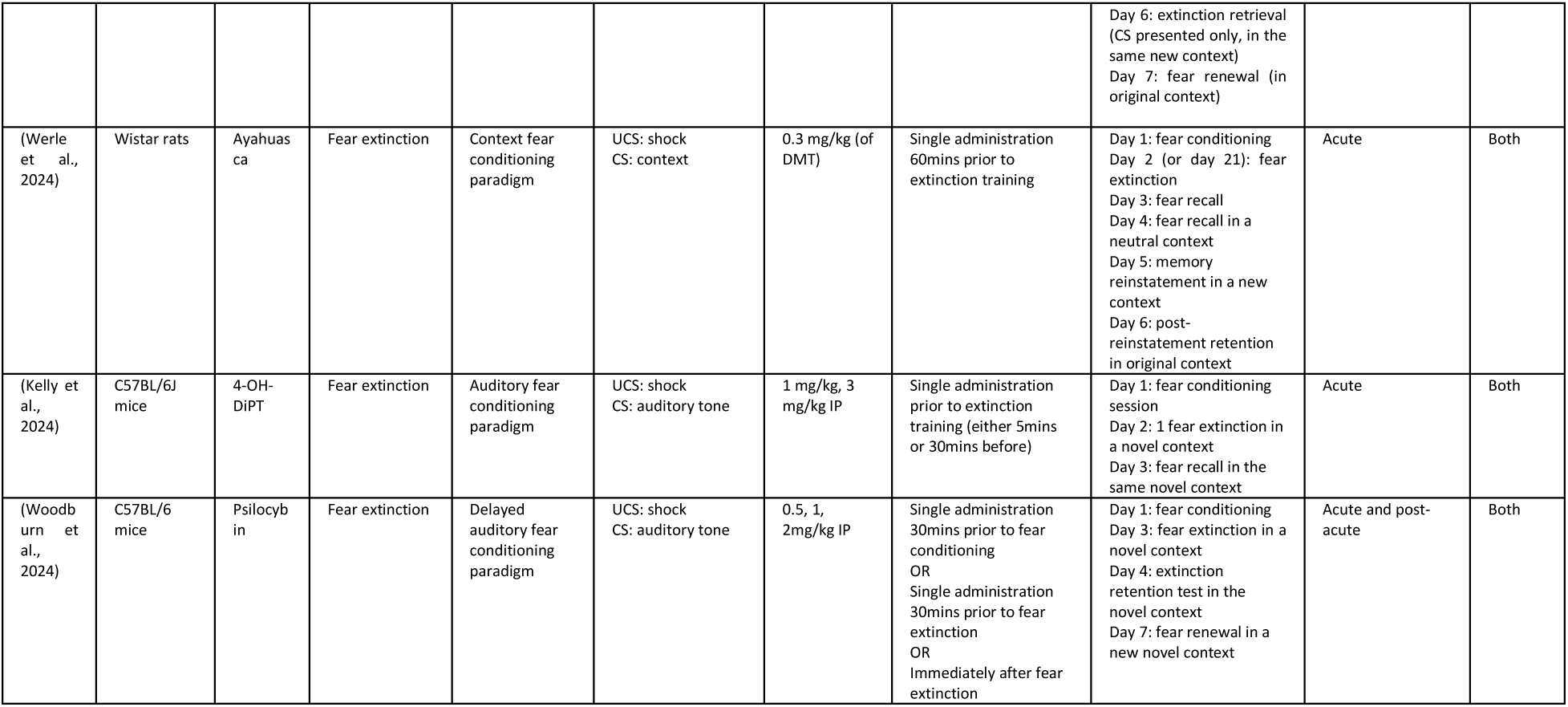
summary of studies investigating the effect of psychedelics on classical conditioning acquisition and extinction. NMR = nictitating membrane response; JMR = jaw movement response; UCS = unconditioned stimulus; CS = conditioned stimulus; IV = intravenous; IP = intraperitoneal; 4-OH-DiPT (4-hydroxy-diisopropyltryptamine), a psilocybin analogue

#### Acquisition of classically conditioned response

Most studies examining the effect of psychedelics on classical conditioning used an aversive unconditioned stimulus, eliciting a defensive response (see Table 1 for a methodological summary).

#### Aversive acquisition conditioning

Most studies used the rabbit nictitating membrane response (NMR) to examine the effect of LSD on classical conditioning acquisition. The NMR is a blink reflex in rabbits, where the nictitating membrane (a third eyelid) contracts in response to an aversive US, such as an air puff or shock, with the learning being the conditioned association with a CS, such as an auditory tone or light. All studies investigated the effect of LSD on the rate at which rabbits acquire a UR (here the NMR) to a CS. Studies varied in the timing of drug administration relative to learning tasks, training length, paradigm-specific features such as the cues used, and the reflex studied.

LSD (compared to vehicle), administered daily, acutely before conditioning sessions, significantly enhanced CS-UCS pairing (Harvey et al., 1988; Schindler et al., 1986; Gormezano and Harvey, 1980; Romano et al., 2010; Welsh et al., 1998; Gormezano et al., 1980). Five studies examined the effect of LSD on the NMR (Gormezano and Harvey, 1980; Romano et al., 2010; Welsh et al., 1998; Schindler et al., 1986; Harvey et al., 1988) and LSD was also found to enhance conditioning of the jaw reflex, an appetitive reflex, in one study (Gormezano et al., 1980). Training schedules typically consisted of daily sessions, where the aversive stimulus was repeatedly presented with the neutral stimulus. Training took place daily for 10 days (Gormezano and Harvey, 1980; Gormezano et al., 1980; Harvey et al., 2004; Schindler et al., 1986), or 8 days (Romano et al., 2010), and one study investigated training for 5 or 10 days with equal numbers of stimulus pairings overall (Welsh et al., 1998). Unfortunately, the effect of training intensity vs length was not analysed in this study.

In terms of stimuli, corneal air puffs were used in two studies (Welsh et al., 1998; Romano et al., 2010), electric shocks were used in three (Gormezano and Harvey, 1980; Harvey et al., 1988; Schindler et al., 1986), and another used an appetitive stimulus (water) (Gormezano et al., 1980). Auditory conditioned cues were used in two studies (Romano et al., 2010; Harvey et al., 1988) and both visual and auditory in four (Welsh et al., 1998; Schindler et al., 1986; Gormezano and Harvey, 1980; Gormezano et al., 1980). There was no clear difference in acquisition between the different stimuli, suggesting that the enhancement of conditioning is not dependent on the cue used.

Most studies used 30nmol/kg LSD with IV administration compared with vehicle (Harvey et al., 1988; Schindler et al., 1986; Gormezano and Harvey, 1980; Welsh et al., 1998; Gormezano et al., 1980) meaning it was not possible to examine dose-response relationships. The only study which used variable doses of LSD (doses 1, 3, 10, or 30nmol/kg), administered it intrahippocampally 20 minutes prior to each conditioning session. A small but significant dose-dependent acceleration of learning was observed in the LSD groups (for all doses other than the lowest) (Romano et al., 2010). An interesting temporal relationship was observed, with smaller doses corresponding with earlier accelerations in learning, and higher doses with later accelerations.

To investigate underlying mechanisms for the increased classical conditioning, the effect of LSD on sensitivity to both the conditioned and unconditioned stimuli was examined to understand whether LSD may be altering environmental or perceptual sensitivity. Two studies showed no difference in acquisition for varying US (shock) intensities; however, an enhancement of acquisition was observed for a greater range of tone (CS) intensities, with a lower acquisition threshold (Gormezano and Harvey, 1980; Gormezano et al., 1980). This is suggestive of enhanced CS sensory processing driving enhanced classical conditioning. In contrast, another study found that LSD increased the amplitude of the NMR itself, with a lower shock threshold required to elicit this response, suggestive of an enhanced sensitivity to the shock (US) as well as the tone (Harvey et al., 1988).

One study examined the effect of LSD on the maintenance and extinction of the conditioned response (Schindler et al., 1986). Again, enhanced acquisition of CRs to both tone and light CS was observed. Maintenance was tested for four days after conditioning with a switch in drug administration leaving four groups (LSD-LSD, vehicle-vehicle, LSD-vehicle, vehicle-LSD). Evidence of state-dependent learning was observed, with a maintenance in performance when the LSD group continued on LSD, and a drop in performance when the LSD group were switched to vehicle, implying that LSD-induced learning did not transfer to the drug-free state. When examining extinction, CRs acquired under LSD were more rapidly extinguished when LSD was continued during extinction training, but this was not the case when CRs were acquired after vehicle. This may reflect an enhanced adaptation to new contingencies under LSD, or alternatively, that material learnt under LSD is more vulnerable to extinction.

There was limited investigation into the underlying biological mechanisms. Ritanserin (5-HT_2A/C_ antagonist) dose-dependently blocked LSD-mediated enhancement in the acquisition of the CR, suggesting that these receptors are required for this enhancement in associative learning (Welsh et al., 1998). Chronic LSD administration resulted in dose-dependent desensitisation of the 5-HT_2A_ receptor, as measured by a decrease in head bobs after post-study administration of DOI (Romano et al., 2010), which the authors speculate may explain the early acceleration and plateau in performance under lower dose LSD. However, this does not explain the later improvements observed in the high-dose LSD group, unless higher-dose LSD is driving performance via other mechanisms.

In summary, LSD appears to increase the acquisition of classically conditioned responses; however, several limitations were noted, including testing in a highly selective population (rabbits), being limited to only two reflexes, and while replicated several times, all studies appeared to come from the same research group. The repeated administration paradigms introduce uncertainty as to whether the observed effects are driven by acute learning changes under LSD, or whether they relate to post-acute cognitive or neuroplastic processes which may underwrite improved learning. Both hypotheses are supported by various results: that an enhancement was seen in some studies as early as the first conditioning session (Gormezano and Harvey, 1980; Gormezano et al., 1980), and that learning was also enhanced later in training (Romano et al., 2010). Furthermore, the observed state-dependent learning suggests that material learnt while under LSD may not translate well to the non-drug state, which has clear implications for psychedelic use in humans.

### Appetitive acquisition conditioning

The research on appetitive conditioning is more limited. One study investigated the effect of LSD on conditioning of the jaw movement reflex (JMR) in rabbits in response to a water reward, finding an enhancement, consistent with results from aversive conditioning paradigms (Gormezano et al., 1980). Two newer studies examined the effect of psychedelics on appetitive classical conditioning, with a primary focus on underlying mechanisms (Martin et al., 2024; Nardou et al., 2023).

One study used a Pavlovian conditioning paradigm, where reward delivery was paired with various cues (Martin et al., 2024). This study has been classified as a classical conditioning study because although it involves reward delivery, the primary measurement was dopamine (DA) transients to reward predictors, which can be considered an unconditioned response. The effect of DOI on nucleus accumbens (NAcc) phasic DA release (signalling prediction error) was measured using fibre optometry. After confirming that DA transients shifted during learning from the cue predicting immediate reward, to an earlier predicting cue (as typically occurs during learning, (Schultz et al., 1997)), it was shown that DOI disrupted this effect. Dose-dependent increases in DA transients were observed to the cue predicting immediate reward, with no change to the earlier cue. Importantly, this was independent of changes in reward value.

Another study, while not purely testing classical conditioning, used a classical conditioning paradigm (social conditioned place preference (sCPP)), to show that psychedelics reopen a critical period for social reward learning in adult mice (Nardou et al., 2023). For the sCPP paradigm adult mice were conditioned to associate one type of bedding with socialising, and another with isolation. Psychedelic administration (ranging from 48 hours to 4 weeks before the task) led to a renewed preference for the social-associated environment. Notably, this behaviour is typically restricted to juvenile developmental windows. The behaviour was observed for weeks after psychedelic administration and was proportional to the length of subjective effects in humans for each drug tested. The paradigm design was such that the ‘test’ followed the isolation conditioning, raising doubt about whether the drugs may have been altering some other behavioural parameter such as an increased desire for novelty, an enhanced sense of boredom, or perhaps an altered sensitivity to rewards. However, the paradigm was also tested in juvenile mice (using MDMA), and there was no increase in the magnitude of social reward learning in this group. Using ex-vivo whole-cell voltage-clamp recordings, the critical period was found to be underwritten by oxytocin-mediated long-term depression (LTD) in the nucleus accumbens, accompanied by transcriptional changes in extracellular matrix remodelling genes.

These studies provide plausible mechanisms whereby an enhancement of learning may be driven by an enhanced environmental sensitivity, by amplifying reward-prediction errors, or regression to a juvenile learning state (which itself would be consistent with an enhanced sense of novelty).

#### Extinction of classically conditioned response

Seven studies examined the effect of psychedelics on fear extinction (for an explanation of fear extinction, see Box 2). Study designs differed in psychedelic used, dose, paradigm, and timings between drug administration and learning, and are summarised in Table 1. Three studies used psilocybin, one used a psilocybin analogue, two used ayahuasca, and one used DOI.

Five studies demonstrated enhanced fear extinction following a single acute administration of a psychedelic within 1 hour before extinction training (Woodburn et al., 2024; Kelly et al., 2024; Werle et al., 2024; Du et al., 2023; Pędzich et al., 2022). Precise drug administration timings before extinction training varied. One study compared 4-OH-DiPT administration 5 minutes vs 30 minutes before, with greater extinction observed for the 5min group (Kelly et al., 2024).

Three studies administered the drug before fear conditioning itself (Woodburn et al., 2024; Catlow et al., 2013; Favaro et al., 2015). Psilocybin administered 30 minutes prior revealed no change in acquisition or subsequent extinction (Woodburn et al., 2024). Psilocybin administration 48 hours prior revealed no effect on fear acquisition, but revealed enhanced fear extinction, only for lower doses (0.1-0.5mg/kg) (Catlow et al., 2013). 30 days of ayahuasca, stopping 48 hours before fear conditioning, showed increased conditioned fear responses. This was only observed for context-cues, and not tone-conditioned cues, and for the lowest dose (120mg/kg) (Favaro et al., 2015).

Heterogeneity in doses, routes of administration, and variable timings between drug administration and paradigms makes interpretation of dose effects challenging. Five studies examined variable doses on fear extinction. IP Psilocybin (0.5mg/kg, 1mg/kg and 2mg/kg) 30 minutes before extinction enhanced it at all doses tested (Woodburn et al., 2024). Dose-dependency was observed for acute IP psilocybin (0.1, 0.5 and 2.5mg/kg) (Du et al., 2023), and acute IP 4-OH-DiPT (Kelly et al., 2024). IP Psilocybin (0.1, 0.5mg/kg, 1mg and 1.5 mg/kg) administered 48 hours before conditioning had no effect on acquisition, however 0.1 and 0.5mg/kg enhanced fear extinction (Catlow et al., 2013). That lower doses enhanced fear extinction is particularly surprising given the early administration. Only the lower dose of oral ayahuasca (120mg/kg) was found to enhance fear conditioning when administered chronically up until 48 hours before conditioning began (Favaro et al., 2015).

Four studies examined the effect of a psychedelic on extinction retention. Two psilocybin studies showed sustained fear extinction on testing 1 and 2 days later (Woodburn et al., 2024; Du et al., 2023). One showed enhanced retention for all doses tested (0.1, 0.5 and 2.5mg/kg) (Du et al., 2023), the other showed an effect for 1mg/kg, but not for 0.5 or 2mg/kg (Woodburn et al., 2024). Both showed sustained suppression of fear renewal a week later, in the original context, and a new context, respectively (Du et al., 2023; Woodburn et al., 2024), implying that this may be retained in novel environments. Contrastingly, fear extinction was not maintained the day after ayahuasca (Werle et al., 2024), DOI (Pędzich et al., 2022), or 4-OH-DiP (Kelly et al., 2024). Retention of learnt associations was modulated by training intensity or dose. When shock strength (duration and amplitude) was increased, extinction was maintained the following day (Pędzich et al., 2022). When an extinction paradigm was repeated twice (including two ayahuasca administrations), extinction learning was enhanced, and was maintained the following day (Werle et al., 2024). Freezing behaviour was reduced after fear reinstatement, suggesting that ayahuasca may be protective against spontaneous recovery of fear memories (Werle et al., 2024).

In terms of sex differences, four studies used male rodents only (Favaro et al., 2015; Pędzich et al., 2022; Du et al., 2023; Catlow et al., 2013). No sex differences were observed in fear extinction after ayahuasca (Werle et al., 2024). Sex specific effects were observed in two studies. Females exhibited a greater reduction in fear extinction, and a sustained reduction in avoidance behaviour in the week following 4-OH-DiP (Kelly et al., 2024). Females responded to a narrower range of doses of psilocybin, with 1mg/kg being optimal, however quantitative analysis was not performed (Woodburn et al., 2024).

Underlying mechanisms were explored in all but one study. Four studies investigated the role of serotonin receptors in psychedelic-induced modulation of fear extinction and recall. Infralimbic 5-HT_2A_ antagonism diminished the enhancing effect of ayahuasca on extinction learning, while 5-HT_1A_ antagonism selectively reduced improvements in extinction recall (Werle et al., 2024). Similarly, 5-HT_2A_ receptor antagonism abolished psilocybin’s effects on extinction, retention, and renewal, while 5-HT_1A_ antagonism affected renewal only (Woodburn et al., 2024), indicating a differential role of these receptors in fear learning processes. DOI had no effect on fear expression in a 5-HT_2A_ knockout model, unlike wildtype mice (Pędzich et al., 2022). Ketanserin (non-selective 5-HT antagonist) reduced the effect of 4-OH-DiPT on fear extinction (Kelly et al., 2024). 4-OH-DiPT promoted spontaneous inhibitory postsynaptic currents in the basolateral amygdala (BLA), revealing a plausible mechanism for psychedelic-induced fear extinction, where 5-HT_2A_ activation on GABAergic interneurons leads to increased BLA inhibition (Kelly et al., 2024), a key area for extinction-induced plasticity (Sierra-Mercado et al., 2011). DOI injected into the amygdala suppressed fear expression, and this was reversed by 5-HT_2A_ antagonism, however no effect was observed for retention (Pędzich et al., 2022).

Contextual fear conditioning is considered a hippocampal-dependent learning task (for review, see (Lopresto et al., 2016). Two studies investigated the effect of psychedelics on hippocampal plasticity. A dose-dependent decrease in BrdU+ neural progenitor cell survival was observed 14 days after psilocybin administration (and this effect was blocked by ketanserin). However, a non-significant increase was observed after the lowest dose (0.1mg/kg), the same dose at which behavioural changes were observed (Catlow et al., 2013). Psilocybin (2.5mg/kg) reversed several deficits induced by fear conditioning 7 days after administration, including structural (hippocampal dendritic complexity and spine density), protein (BDNF and mTOR) and neurogenesis (DCX+ and BrdU+ cells in the hippocampal dentate gyrus) (Du et al., 2023).

In summary, it appears that psychedelics, particularly when administered acutely before extinction training, can facilitate fear extinction in animal models, although the durability of this effect varies. Dose, timing, training intensity, and sex differences substantially modulate outcomes, and the results for fear renewal in novel contexts are mixed. Identified mechanisms appear to relate to serotonin receptor modulation.

### Operant conditioning

#### Box 3: operant conditioning

Operant learning is a form of associative learning in which the probability of a behaviour is modified by its consequences, where actions are reinforced when followed by rewards and diminished when followed by punishments or the absence of reinforcement. Unlike classical conditioning, where associations form between stimuli, operant learning is driven by voluntary behaviours and their outcomes.

##### Negative reinforcement

All studies examining negative reinforcement utilised avoidance learning paradigms, where animals learn to proactively avoid an aversive stimulus after forming an association with a predictive cue. Most studies used the shuttle box, a behavioural apparatus containing two compartments which the animal can move between. Following conditioning to associate an auditory tone (CS) with a shock (US), the animal can flee in response to the tone, known as the conditioned avoidance response (CAR). Six studies examined the effect of LSD on the CAR, by measuring the latency from CS presentation to the CAR, or the percentage of successful CARs.

##### Positive reinforcement

Only one study examined positive reinforcement and used a Skinner box lever-pressing paradigm. In this paradigm, animals are placed in a chamber equipped with a lever, which they learn to press to receive a reward, such as a food pellet.

##### Reversal learning

Reversal learning is a form of flexible learning that involves the ability to modify behaviour in response to changing reinforcement contingencies (for review, see (Izquierdo et al., 2017)). It requires the suppression of previously learnt associations and the acquisition of new stimulus-outcome or action-outcome associations. Reversal learning paradigms typically involve associating stimuli with rewards or punishments (which may be probabilistic themselves) before a rule change where these contingencies are reversed, known as the ‘reversal’.

Sixteen studies examined the effects of psychedelics on operant learning (see Box 3). Study details are summarised in Table 2.

**Table 2:** summary of studies investigating the influence of psychedelics on operant learning. SC=subcutaneous, IV=intravenous, IP=intraperitoneal, CAR = conditioned avoidance response

| Reference | population | psychedelic | Learning mechanism | Paradigm | Stimuli used | dose | Frequency and timing of drug administration | Number of training sessions | Acute or post-acute effects | %female |
| --- | --- | --- | --- | --- | --- | --- | --- | --- | --- | --- |
| (Bignami, 1972) | Wistar albino rats | LSD | Negative reinforcement | Shuttle box - Escape and avoidance learning | UCS: shock<br>CS: light<br>CR: avoidance<br>Measurement: % CAR | 0.05, 0.2 mg/kg SC | 30mins before each session | Daily sessions of avoidance training for 6 days | Unclear – acute and post-acute due to repeated administrations | All male |
| (Stoff et al., 1974) | Hooded rats | LSD | Negative reinforcement | Shuttle box - Avoidance learning | UCS: shock<br>CS: light and auditory tone<br>CR: avoidance<br>Measurement: latency from CS to CAR | 0.1, 0.5mg/kg IP | 10mins before acquisition test + daily administration (0.5mg/kg) for the preceding 4 days<br>OR<br>24 hours before initial acquisition test | Day 1: initial acquisition test<br>Day 2: second acquisition test | Acute and post-acute | All male |
| (Izquierdo, 1975) | Wistar rats | LSD | Negative reinforcement | Shuttle box - Habituation, pseudoconditioning and avoidance learning | UCS: shock<br>CS: auditory tone<br>CR: avoidance<br>Measurement: % CAR | 0.075, 0.3 mg/kg IP | 5 mins before first session<br>OR<br>Immediately after first session. | Day 1: habituation or pseudoconditioning or avoidance learning session<br>Day 8: retention session | Acute | All female |
| (Waser et al., 1976) | Wistar rats | LSD | Negative reinforcement | Y-maze task - Avoidance learning | UCS: shock<br>CS: light<br>CR: avoidance<br>Measurement: % CAR | 5, 20, 100 µg/kg IP | 30mins before each session | Day 1-23: training twice per week with LSD<br>Day 24: training without LSD<br>Day 25-28: training twice per week with LSD<br>Day 29-32: training twice per week without LSD | Unclear – acute and post-acute due to repeated administrations | All male |
| (Castellano, 1979) | C57BL/6 mice | LSD | Negative reinforcement | Shuttle box - Escape and avoidance learning | UCS: shock<br>CS: light<br>CR: avoidance<br>Measurement: % CAR | 0.05, 0.1, 0.2 mg/kg IP (pre-trial).<br>0.2 mg/kg IP (post-trial) | 15mins before each session<br>OR<br>Immediately, or 2 hours after each training session. | Daily avoidance training for 5 days. | Unclear – acute and post-acute due to repeated administrations | All male |
| (Buchborn et al., 2014) | Wistar rats, including rat model of depression (olfactory bulbectomy (OBX)) with sham-operated rats as controls (sham). | LSD | Negative reinforcement | Pole jumping - One-way active avoidance learning | UCS: shock<br>CS: auditory tone<br>Measurement: % CAR | 0.13 mg/kg SC | 6 days of LSD administration without training, followed by 5 days of LSD administration 2 hours after training sessions. | Daily training for 5 days.<br>Factorial design with 4 groups: sham/saline vs sham/LSD; OBX/saline vs OBX/LSD | Post-acute | All male |
| (Rosen and Buga, 1973) | Albino rats | LSD | Positive reinforcement | Skinner box lever press OR runway task for food reward | Action: lever press or alley run<br>Outcome: food reward<br>Measurement: differential response speed (for positive or negative bar) | 0.05, 0.20 mg/kg IP | Administration 25mins before lever testing sessions.<br>OR<br>10 mins before alley runs. | Lever training for 3 weeks. 5 testing sessions every 72 hours with subsequent further testing with factorial design for drug administration.<br>Runway task – training for 3 weeks. Testing once weekly for 2 weeks. | Unclear – acute and post-acute due to repeated administrations | All male |
| (Butters, 1966) | Albino rats | LSD | Reversal learning (perseveration) | T-maze discrimination reversal learning task | Cues: position (L/R doorways), or brightness (light/dark doorways)<br>Incentive: shock<br>Measurement: number of perseverative responses, and trials to criterion | 0.5mg/kg IP | Administration 8 mins before contingency change | Position training followed by brightness training and vice versa. | Acute | Unclear |
| (King et al., 1974) | Wistar SPF rats | LSD | Reversal learning | brightness discrimination reversal learning task. | Cues: brightness (light/dark tunnels)<br>Incentive: food reward<br>Measurement: p(correct response) after reversal | 6.25, 12.5, 25, 50 µg/kg IP | 5, 15, 45 or 90mins before reversal | Day 1-8: daily acquisition training sessions<br>Day 9-11: rest<br>Day 12: reversal session | Acute (LSD half-life in rat estimated at 20 mins (Freedman, 1969)) | All male |
| (Kanen et al., 2023) | Humans | LSD | Reversal learning | Probabilistic reversal learning task | Cues: Three abstract visual stimuli<br>Incentive: green smiling face for reward, red frowning face for punishment.<br>Measurement: win-stay/ lose-shift, perseverative errors | 75µg IV | 5 hours before task | 2 task administrations per participant (cross-over design) | Acute | All male |
| (Torrado Pacheco et al., 2023) | Long-Evans rats | Psilocybin | Reversal learning | Reversal learning task | Cues: auditory tone or light<br>Incentive: food reward or shock<br>Measurement: R-value (cue vs baseline port entries) | 1mg/kg IP | 20 mins before reversal | Training over 10 sessions.<br>Contingencies reversed on 6<sup>th</sup> session. | Acute | Both |
| (Šabano<br>vić et<br>al.,<br>2024) | C57BL/6J mice | DOI | Reversal<br>learning | 2-step<br>probabilistic<br>reversal<br>learning task | Cues: distinct<br>auditory cues for<br>reward or omission<br>Incentive: water<br>Measurement:<br>Speed of reversal<br>adaptation; trial-by-trial learning<br>from reward,<br>omission, and<br>transition x<br>outcome<br>interactions | 2mg/kg IP | 1 week before novel<br>transition reversal | Weeks 1-6: training<br>Week 7: drug<br>administration and<br>continued training<br>Week 8-9: transition<br>reversal and task<br>continuation | Post-acute | Male |
| (Conn et<br>al.,<br>2024) | Sprague-Dawley rats<br>with activity-<br>based anorexia<br>rat model | Psilocybin | Reversal<br>learning | Reversal<br>learning task | Cues: nose-poke<br>ports with FR5<br>training schedule<br>Incentive: food<br>reward<br>Measurement:<br>Accuracy,<br>perseverative<br>errors, trials to<br>criterion, response<br>latencies, reward<br>efficiency | 1.5 mg/kg IP | 18 hours before<br>reversal phase | Week 1: Up to a week of<br>training (until high<br>accuracy was achieved)<br>drug<br>Week 2: reversal followed<br>by 3 days of testing | Post-acute | All<br>female |
| (Fisher<br>et al.,<br>2024) | Sprague-Dawley rats | Psilocybin | Reversal<br>learning | Modified two-armed bandit<br>reversal<br>learning task | Cues: nose-poke<br>ports<br>Incentive: sucrose<br>Measurement:<br>rewards earned,<br>losses, reversals<br>completed,<br>response vigour | 1.5mg/kg IP | 24 hours before first<br>session | 14 consecutive daily<br>sessions (3hr/day),<br>voluntary engagement | Post-acute | All<br>female |
| (Casano<br>va et al.,<br>2024) | Humans | Psilocybin | Probabilisti<br>c<br>reinforcem<br>ent learning | Probabilistic<br>cue-reward<br>learning task | Cue/action: neutral<br>or fearful faces<br>either conscious<br>(long exposure) or<br>subconscious (brief<br>exposure).<br>Followed by choice<br>between 2 abstract<br>symbols with<br>different reward<br>probabilities (70%<br>vs 30%).<br>Incentive:<br>monetary reward<br>Measurement:<br>accuracy, trial-by-trial behaviour | 10 mg, 15<br>mg, 20 mg<br>route not<br>specified | 3-4 hours before task<br>administration | 2 task administrations per<br>participant (crossover<br>design) | Acute | Both |
| (Jacobs<br>et al.,<br>2024) | Long Evans rats | Psilocybin | Probabilisti<br>c<br>reinforcem<br>ent learning | Probabilistic<br>punishment<br>task | Action: Chained<br>seek–take nose-poke responses in<br>operant chamber<br>Outcome: Seek<br>action<br>probabilistically<br>punished (0–60%<br>foot shock risk);<br>take action always<br>safe and rewarded<br>with sucrose pellet<br>Measurements:<br>trial completion<br>rate, latency,<br>behavioural<br>suppression | 1mg/kg IP | 15 mins before 1<sup>st</sup><br>punishment session<br>OR<br>After punishment<br>learning | Days 1-4: seek-take<br>action training<br>Days 5-12: punishment<br>training where<br>probabilistic foot shock<br>introduced after seek<br>response<br>Day 13: additional<br>sessions | Acute | Both |

### Negative reinforcement

All studies examining negative reinforcement utilised avoidance learning paradigms (see Box 3). Six studies examined the effect of LSD on the conditioned avoidance response (CAR), by measuring the latency from CS presentation to the CAR, or the percentage of successful CARs. Four studies used shuttle box paradigms, one used a Y-maze avoidance learning task, and one used a pole-jumping avoidance task.

Five studies demonstrated an enhanced acquisition of CARs (Bignami, 1972; Stoff et al., 1974; Izquierdo, 1975; Castellano, 1979; Buchborn et al., 2014), and one did not (Waser et al., 1976). There was considerable heterogeneity in methodology, in terms of timings between the drug administration and training sessions, timing and length of training itself, drug doses, and rodent strains.

Three studies repeated training on 5-6 consecutive days with daily LSD administration (Bignami, 1972; Castellano, 1979; Buchborn et al., 2014), and two used 1 administration/acquisition session only (Izquierdo, 1975; Stoff et al., 1974). The rate of learning enhancement differed; enhanced learning only emerged later in training in one study (Bignami, 1972), however in three studies, increased learning was observed immediately (Stoff et al., 1974; Izquierdo, 1975; Castellano, 1979). One study compared the effect of repeated doses of LSD for the five days preceding training, with the effect of a single dose, and there was no observed difference between them (Stoff et al., 1974). One study (Waser et al., 1976) used a Y-maze task to examine changes in avoidance learning over a longer timescale. IP LSD (5, 20, and 100 µg/kg) administered 30 minutes before conditioning sessions, which took place twice a week across several weeks, significantly delayed the acquisition of CAR. LSD-treated rats only caught up with the vehicle group 5–7 sessions later. State-dependent learning was observed, where after LSD withdrawal (at session 24) performance declined significantly, but was restored when LSD was reintroduced.

Dose-related performance improvements were noted in one study (LSD 0.05, 0.1, 0.2mg/kg (Castellano, 1979)), with a partial relationship seen in another (LSD 0.05, 0.1, 0.2mg/kg), where lower doses had a greater effect on learning (Bignami, 1972). No difference was seen between 0.1 vs 0.5mg/kg LSD (Stoff et al., 1974). 0.075mg/kg LSD had a significant effect compared with 0.3mg/kg which had no effect (Izquierdo, 1975).

One study investigated the effect of acute LSD on pseudoconditioning (Izquierdo, 1975). Pseudoconditioning differs from conditioning in that it involves presenting both tones and shocks without contiguity. LSD dose-dependently enhanced avoidance of the tone in pseudoconditioning trials. This raises uncertainty about whether LSD does in fact enhance associative learning, or whether the results are driven by other factors, such as changes in environmental sensitivity or inference. This is indeed supported by the finding that habituation was blocked by LSD, in a dose-dependent manner (Izquierdo, 1975), implying that each presentation may have been endowed with a sense of novelty, or could be explained by an increased environmental sensitivity.

LSD was generally administered shortly before training began (ranging 5-30 minutes), meaning that the animals were under the acute effects while they learnt (with the exception of (Buchborn et al., 2014) and one experiment in (Castellano, 1979)). Immediate LSD administration post daily training, but not 2 hours after, significantly enhanced the CAR, suggesting that this is due to acute, rather than post-acute effects (Castellano, 1979). No change was observed when LSD was administered 24 hours prior to conditioning (Stoff et al., 1974), however no statistics were reported for this result. In contrast, LSD administration 2 hours after each conditioning session led to a significant improvement in CAR, and notably, a mouse model with known avoidance learning deficits was used in this study (Buchborn et al., 2014).

Two studies noted a normalisation effect of LSD, suggesting that worse performers may benefit more (Waser et al., 1976; Buchborn et al., 2014). Olfactory bulbectomised (OBX) mice are an animal model of depression known to exhibit avoidance learning deficits (for review see (Song and Leonard, 2005)). Normalisation of learning following repeated LSD (0.13 mg/kg, sc) administration 2 hours after daily pole-jumping conditioning sessions was observed in OBX mice (Buchborn et al., 2014). No enhancement was observed in the wildtype animal/LSD group. Interestingly, such avoidance learning deficits are reversible by drugs classified as antidepressants (Kelly et al., 1997). Neurochemical analysis revealed that LSD normalised hippocampal 5-HT_2_ receptor signalling in OBX rats, with no effect seen in control animals (Buchborn et al., 2014). These findings suggest that LSD reverses avoidance learning deficits via modulation of serotonergic signalling, in a manner which may be similar to antidepressants.

Three studies tested retention of avoidance learning in a drug-free state. The day after avoidance training under LSD, faster avoidance responses were observed (Stoff et al., 1974). Retention a week following a single acquisition session (with LSD) revealed no difference in retention for the low-dose group (0.075mg/kg), and a worsened effect for the high-dose group (0.3mg/kg) (Izquierdo, 1975). Immediate post-trial LSD had no effect on retention (Izquierdo, 1975). This is in contrast to another study which found a significant improvement in retention when LSD was administered immediately, but not when it was administered 2 hours later (Castellano, 1979).

Various rodent strains were used. Four studies used Wistar rats (Izquierdo, 1975; Bignami, 1972; Waser et al., 1976; Buchborn et al., 2014), one used hooded rats (Stoff et al., 1974) and one used C57BL/6 (C57) mice (Castellano, 1979). One study observed that LSD may improve the performance of poor performers, and have a deleterious effect on good performers (Waser et al., 1976). Castellano describes his strain as ‘high arousal’ and poor experimental performers (Castellano, 1979). This normalisation hypothesis is further supported by the OBX mice doing well (Buchborn et al., 2014).

In summary, there appears to be a consistent relationship between LSD administration and enhanced avoidance learning within shuttle box paradigms. The effects are more robust when LSD is administered shortly before or immediately after training, with mixed results for longer durations between drug administration and training, suggesting enhancements may depend on the acute drug effects. However, whether this is retained and incorporated into sustained behavioural change is less clear. One of the two studies that used a non-shuttle box paradigm demonstrated worse learning (initially), implying that results obtained from a specific paradigm may not be widely generalisable. Substantial methodological variability, including differences in dosing, timing of administration relative to training, duration of training sessions, and rodent strains, makes translation challenging.

### Positive reinforcement

The research on the effects of psychedelics on positive reinforcement is comparatively scarce, with only two studies identified.

One study examined the effect of LSD on instrumental conditioning using a Skinner box lever pressing paradigm. Rats administered a lower dose (0.05mg/kg) of LSD exhibited enhanced reinforcement learning, driven by faster responding to the rewarding bar, and slower learning under a high dose (0.2mg/kg) (Rosen and Buga, 1973), suggesting that lower dose LSD may enhance reward learning. This study also examined learning from omissions, where animals were trained to run a runway for a food reward (Rosen and Buga, 1973), before the reward was removed. In the initial learning phase, acute LSD (0.20 mg/kg) impaired acquisition. Following reward removal, LSD increased resistance to extinction in the runway task, in a dose-dependent fashion, suggestive of increased perseverative behaviour, or alternatively, a decreased sensitivity to negative feedback.

Another study (discussed already in ‘Classical Conditioning’ (Martin et al., 2024)) used a lever press paradigm with food or water rewards, with an escalating fixed ratio (FR) design, meaning that animals had to gradually work harder for their rewards. Acute DOI had differential effects on the reward value itself, with water becoming more rewarding, and food becoming less rewarding. However, increased dopamine signalling to the immediate reward-predicting cue was observed for both reward types, suggesting that this increase in dopamine prediction error signalling is not mediated by changes in reward value. At higher effort requirements, DOI reduced responding for both rewards, suggesting a general decrease in willingness to exert effort.

In summary, studies examining the effects of psychedelics on positive reinforcement are limited and present somewhat mixed results. While low-dose LSD may facilitate the acquisition of instrumental responding for rewards, higher doses impair task performance, suggesting an inverted-U or otherwise complex dose-response relationship, which is likely confounded by acute drug effects. Effects observed, such as enhanced resistance to extinction or impaired responding at higher task demands, may reflect changes in reward sensitivity, perseverative behaviour, or motivation.

### Mixed reinforcement

#### Reversal learning

Seven studies examined the effect of a psychedelic on reversal learning (Box 3); five used deterministic reversal learning paradigms (Torrado Pacheco et al., 2023; King et al., 1974; Butters, 1966; Conn et al., 2024; Fisher et al., 2024), and two used probabilistic reversal learning paradigms (Šabanović et al., 2024; Kanen et al., 2023). See Table 2 for study details, including outcome measurements.

Three studies showed an enhancement of reversal learning (King et al., 1974; Conn et al., 2024; Šabanović et al., 2024) following a psychedelic, one showed an enhancement and reduction, depending on the stimuli used (Butters, 1966), and one showed no change (Torrado Pacheco et al., 2023). Two studies used reinforcement learning models to examine behavioural changes, with more complex outcomes, and these will be discussed separately (Kanen et al., 2023; Fisher et al., 2024).

Four studies examined the effects of drug administration on reversal learning acutely (Torrado Pacheco et al., 2023; King et al., 1974; Butters, 1966; Kanen et al., 2023), and three examined post-acute effects (Conn et al., 2024; Šabanović et al., 2024; Fisher et al., 2024). Timings between drug administration and task reversal varied, with the drug administered usually after the acquisition phase and prior to the reversal. LSD, psilocybin and DOI were all shown to enhance reversal learning, suggesting that this may be a broad effect of psychedelics. Interestingly, where an improvement in reversal learning was observed for LSD, it was not observed with 2-bromo-LSD (BOL-148), a non-psychoactive analogue (King et al., 1974).

Acute LSD (≥12.5 µg/kg) significantly improved reversal learning in rats in a light/dark discrimination task (King et al., 1974). This was observed for drug administration timings prior to the reversal, which ranged from 5 minutes to 90 minutes before. This was not replicated in another study which used a reversal learning paradigm with cues associated with both appetitive (food reward) and aversive (foot shock) outcomes (Torrado Pacheco et al., 2023). Psilocybin 20 minutes before the reversal did not affect reversal learning, for either appetitive or aversive outcome. This study did, however, find that psilocybin improved set-shifting, although the influence of psychedelics on behavioural and cognitive flexibility is beyond the scope of this review.

Using an animal model of anorexia, an improvement in reversal learning after psilocybin administration was observed post-acutely using a paradigm requiring positional (left/right) nose pokes for reward (Conn et al., 2024). Mice were initially trained on a fixed ratio schedule to receive rewards from either exclusively the left or right port (counterbalanced between animals). After training, psilocybin was administered, and 18 hours later the contingencies were reversed. Psilocybin vs vehicle enhanced reversal learning, measured by faster adaptation to the contingency reversal, reduced perseverative errors, and increased task engagement. This animal model is known to exhibit deficits in reversal learning and attentional set-shifting (Huang et al., 2023; Allen et al., 2017), again suggestive of normalisation.

The effects of psychedelics on reversal learning are complex and seem to relate to multiple factors such as administration timings, task structure and baseline performance. One study adds a further layer of complexity, suggesting that psychedelics may enhance reversal learning, but that this is dependent on post-drug training (Šabanović et al., 2024). DOI improved adaptation to a novel transition reversal in a two-step probabilistic reversal learning task when the reversal was introduced a week, but not a day, after drug administration (Šabanović et al., 2024). Furthermore, these improvements were dependent on task experience in the week after drug administration, with a lack of task experience in the ensuing week leading to significantly worse performance. This implies an interaction between the DOI and task experience in the post-acute phase. Interestingly, while DOI also appeared to improve adaptations to serial reversals, this was driven by a post-injection performance decline in the vehicle group exclusively. The pre-post measurements in the DOI group revealed no absolute improvements, which may reflect increased resilience to injection stress in the DOI group, again pointing to a possible role of psychedelics normalising deficient learning. Examination of choice behaviour following wins and losses revealed enhanced learning from reward omissions post DOI, with symmetric inference learning. This is a unique behavioural strategy not typically observed in mice which are normally driven by reward learning. An interpretation is DOI conferred an enhanced sensitivity to the environment, with previously overlooked cues now being incorporated into decision making.

Several studies examined the neurobiological effects underlying changes in reversal learning with variable approaches. Biochemical analysis indicated that LSD, but not BOL-148 (a non-psychoactive LSD analogue), increased brain 5-hydroxytryptamine (5-HT) levels 45 minutes post-injection, with no significant effects on noradrenaline or dopamine, suggesting that LSD increased brain 5-HT levels, although this study did not investigate a causal relationship with reversal learning (King et al., 1974). Psilocybin-induced improvements in reversal learning were blocked by 5-HT_1A_, but not 5-HT_2A_ receptor antagonism (Conn et al., 2024). Ex-vivo MRI showed increased volumes in sensory and associative areas on the day following DOI administration, suggestive of rapid changes in structural plasticity (Šabanović et al., 2024). However, there was a temporal disassociation between these structural changes and cognitive changes, which only emerged a week later. As structural changes were not compared between the group which received task experience and the group which had no task experience, it is not possible to infer directionality between learning and neuroplastic changes.

In summary, psychedelics may enhance reversal learning, though effects vary considerably depending on methodological factors such as timing, task structure, baseline performance, and drug-specific mechanisms. Psychedelics may exert these effects by normalising deficient learning, or enhancing sensitivity to previously overlooked environmental cues, likely mediated through serotonergic modulation and neuroplastic changes.

#### Computational modelling of reversal learning

Computational modelling permits a richer characterisation of learning and decision-making. For example, reinforcement learning (RL) models describe how individuals incorporate feedback and uncertainty into learning strategies, using parameters such as the learning rate (how individuals update value representations in response to feedback), reward sensitivity and exploration parameters (Sutton and Barto, 1998). Two studies performed computational modelling to characterise learning changes, one in humans and one in mice.

To understand the effect of LSD on reinforcement learning in humans, a probabilistic reversal learning (PRL) task was completed during the acute drug phase (Kanen et al., 2023). Whereas most of the previous paradigms involved drug administration prior to the reversal and after acquisition, this paradigm involved acquisition and reversal in one session. LSD increased the learning rate (LR) for both reward and punishment outcomes (and especially for rewards). In other words, LSD accelerated the updating of value representations, following a prediction error, especially for rewarding outcomes, conferring a greater sensitivity to reward prediction errors. Interestingly, while the reward LR was increased in both the acquisition and reversal phases, the punishment LR was increased only in the reversal phase, meaning that learning from negative feedback was only enhanced when the environment changed. Strikingly, a similar pattern was also observed in mice (Šabanović et al., 2024), where an increased learning from omissions was observed in the reversal phase only, however given methodological and species difference between studies, this link is made tentatively. Interestingly, under LSD, fewer errors in the acquisition phase predicted more perseverative errors in the reversal phase, suggesting that better initial learning resulted in greater subsequent perseveration. The authors speculate that this may relate to stronger learning under LSD, which is subsequently harder to shift (Kanen et al., 2023). However, whether these enhancements in learning are observed in the post-acute phase are yet to be tested in humans.

One study examined the effect of psilocybin on reversal learning in rats post-acutely, using a two-armed bandit reversal learning task, where psilocybin was administered 24 hours before task onset (Fisher et al., 2024). Task-engagement was optional for 3 hours each day for 2 weeks. While this study primarily measured task engagement, active inference (AI) modelling was used to characterise learning behaviour. Psilocybin increased the forgetting rates for rewards and reduced the forgetting rate for losses. In other words, when the psilocybin-treated rats received a reward, their prior belief was down-weighted compared with the control group. Conversely, the forgetting rate for losses decreased, meaning that experiencing a loss had a reduced influence on updating their priors. While this pattern was also observed in the control group, it was significantly enhanced in the psilocybin group. This is consistent with an optimism bias, where mice learn more from positive than negative outcomes, which may also explain increased task engagement.

Differences in the computational frameworks (reinforcement learning vs active inference) complicate direct comparisons between these studies. Whereas Kanen et al compared only RL models in model comparison, Fisher et al compared RL and AI models, with an AI best explaining the effect of psilocybin on choice behaviour, suggesting that AI modelling in humans may be a useful avenue for future research.

### Probabilistic reinforcement learning

One study, conducted in healthy volunteers, examined the effect of psilocybin on the influence of conscious and subconscious cues (fearful or neutral faces presented before the stimulus) on probabilistic reward learning (Casanova et al., 2024). Overall, acute psilocybin revealed no difference in reward learning vs placebo (measured by accuracy), however, this was driven by better learning in the 20mg group and worse learning in the 15mg group (vs placebo). Learning was impaired in the psilocybin vs placebo group when subconscious cues were presented. The authors speculate that a possible reduction in top-down processing under psychedelics leads to a reduction in automatic suppression of task-irrelevant stimuli, leading to a disruption in task performance. However, this interpretation does not account for why subconscious neutral but not subconscious fearful stimuli disrupted learning under psilocybin (Casanova et al., 2024). While there was a slight increase in switching behaviour under psilocybin, which they suggest may reflect higher exploratory behaviour, this was borderline significant. This is in agreement with the decreased stimulus stickiness parameter observed under LSD (Kanen et al., 2023).

The influence of psilocybin on probabilistic punishment learning was investigated using a chained operant reinforcement paradigm involving an escalating risk of foot-shock during reward-seeking actions (Jacobs et al., 2024). The effect of psilocybin depended on the timing and the sex of the animal. When administered 15 minutes before initial punishment learning, punishment association learning was enhanced selectively in females (measured by increased behavioural suppression to punishment-associated actions). However, when given after learning, psilocybin reduced behavioural suppression in both sexes (meaning that rats became more likely to perform the risky action despite the threat of punishment), suggesting increased resistance to punishment. Psilocybin did not alter sensitivity to the shocks themselves, suggesting that this was not due to changes in pain perception. These findings indicate sex-specific effects of psilocybin on punishment learning. Notably, these effects were not observed with DOI, a 5-HT_2A_ receptor agonist.

## Discussion

This systematic review synthesised studies examining the effect of psychedelics on associative learning mechanisms. The evidence suggests that psychedelics may enhance classical conditioning, fear extinction, avoidance learning, and reversal learning; however, this is largely restricted to animal models. Mechanistically, these learning changes may be accompanied by an increased environmental sensitivity and increased sensitivity to prediction errors. The influence of psychedelics on associative learning may be conceptualised in terms of both the <u>content</u> of the associations formed and the <u>process</u> by which they are acquired. With respect to content, the studies reviewed indicate that psychedelics enhance associative learning for a range of stimuli. However, whether this generalises beyond the paradigms reviewed here, especially to complex human decision making, has yet to be tested. Enhancements were observed in classical, operant, and reversal paradigms, suggesting that this may be a general effect.

Various mechanisms were identified regarding how associative learning may be enhanced. There may be a normalisation effect of psychedelics, with some studies suggesting that only animals with learning deficits improve under psychedelics. At a receptor level, 5-HT_2A_ and 5-HT_1A_ receptors appear to be important, given the results of knockouts and antagonist studies (Pędzich et al., 2022; Welsh et al., 1998; Kelly et al., 2024). This is in line with findings showing a crucial role of serotonin modulation in associative learning, with 5-HT_2A_ receptors on thalamocortical synapses essential for associative learning (Barre et al., 2016). Serotonin modulation also appears highly relevant to reversal learning specifically, with changes in reversal learning observed after pharmacological modulation of 5-HT_2C/1A_ receptors, driven by altered sensitivity to positive and negative feedback (Phillips et al., 2018), and a reduction in cognitive flexibility following prefrontal serotonin depletion (Clarke et al., 2004).

Associative learning theories posit that learning is driven by prediction errors (PE), which are generated by unexpected outcomes, or unexpected omissions, and the size of the prediction error determines the magnitude of learning (Rescorla, 1972). Such error signals have been observed in dopamine neurons during reward learning in primates (Schultz et al., 1997; Schultz et al., 2000) and in humans during Bayesian belief updating from priors to posteriors (Nour et al., 2018). As learning progresses, the dopamine neuron signals shift from firing to the unexpected outcome to the conditioned (predictive) stimulus. The finding that DOI leads to increased PE signals and dopamine transients for fully predictable rewards, akin to those seen in earlier learning stages, suggests that PE signals are present for well-established associations (Martin et al., 2024). This may be explained by a reduction in prior expectation, consistent with the REBUS model (Carhart-Harris and Friston, 2019). Alternatively, it may represent an amplified prediction error itself, perhaps due to an enhanced sense of novelty, consistent with the recently proposed Synthetic Surprise Model (De Filippo and Schmitz, 2024), or environmental salience. An enhanced PE is consistent with the observation that psychedelics enhance the learning rate in humans using reinforcement learning modelling (Kanen et al., 2023). A higher learning rate corresponds with a higher precision of the prediction error (Hohwy, 2017), which implies either a reduction in top-down expectation, or a greater influence of bottom-up information.

While these two studies (Kanen et al., 2023; Martin et al., 2024) examined the acute effects of a psychedelic, one study examined the post-acute effects in rodents (Fisher et al., 2024). Active inference modelling revealed that forgetting rates were enhanced for rewards, consistent with down-weighting of priors for rewards, which again is suggestive of psychedelics enhancing belief updating, in this case in a manner which is consistent with an optimism bias. While this remains to be tested in humans, it implies that this enhancement of prediction error signalling may outlast the acute effects. Characterising this further will have important implications for understanding how learning may be enhanced in the post-acute phase. Furthermore, replicating changes in learning model parameters from animal models in humans will clarify translatability of animal models to humans.

If prediction errors are enhanced under psychedelics, this may explain a heightened sensitivity to the environment, which was suggested by this review. Studies showed an increased sensitivity to either conditioned or unconditioned stimuli in classical conditioning paradigms (Gormezano and Harvey, 1980; Gormezano et al., 1980; Harvey et al., 1988). Fear extinction was reliably enhanced under psychedelics, which is known to be highly context sensitive. Furthermore, mice adopted an unusual behavioural strategy where they learnt from reward omissions, which were previously overlooked, again suggestive of enhanced environmental sensitivity (Šabanović et al., 2024). Crucially, this was observed in the post-acute phase, implying that this sensitivity may outlast the acute effects. An enhanced sensitivity to the environment could be explained by a number of mechanisms, including an enhanced sense of novelty, increased prediction errors, higher learning rates, and in turn reduced priors.

As well as being a potential driver of associative learning, it is possible that this environmental sensitivity could explain the context-dependent learning observed. In this review we reported evidence of state-dependent learning (Schindler et al., 1986; Waser et al., 1976), where learning under LSD was not translated to a drug-free state, but upon re-administration of LSD, this was recovered. In this situation, the drug experience can be considered the ‘context’. One study did show translation of learning (fear extinction) to a novel environment (Woodburn et al., 2024). Context-dependent effects of psychedelics have been observed in humans, for example, the influence of natural surroundings during a psychedelic experience predicted later nature-relatedness (Kettner et al., 2019), however, whether changes in associative learning under a psychedelic translate to a drug-free state in humans remains to be tested. Characterising this will enhance our understanding of how psychedelic insights may be incorporated into individuals’ lives.

Of critical relevance to humans and the psychedelic-assisted psychotherapy model is whether changes in learning outlast the acute effects. The earlier literature reviewed here was dominated by measuring learning under the acute effects of a psychedelic, often with repeated administration regimes, rendering interpretation of acute vs post-acute effects challenging. The post-acute period has been examined more recently, with mechanisms identified including the reopening of critical periods (Nardou et al., 2023), increased avoidance learning in a deficit model (Buchborn et al., 2014), and enhanced fear extinction (Catlow et al., 2013), fear conditioning (Favaro et al., 2015) and reversal learning (Conn et al., 2024; Fisher et al., 2024; Šabanović et al., 2024). For reversal learning, it is noteworthy that an interaction between the environment and the drug (DOI) was observed, meaning that only animals who were exposed to the task in the week after the drug showed any improvement in reversal learning. This suggests that behavioural changes may not be immediate, and may in fact develop during a window of enhanced learning. This is supported by the finding that there is a critical window of plasticity after a range of psychedelic drugs (Nardou et al., 2023). It therefore seems possible that psychedelics may induce complementary mechanisms whereby an enhancement in learning is underwritten by a period of neuroplasticity, providing a catalyst to change behaviour.

Returning to the ‘content’ and ‘process’ of how psychedelics change associative learning, the present review suggests that associative learning may be enhanced by a range of psychedelics, and the limited but accumulating evidence suggests that this may be driven by changes to prediction error learning. The evidence also suggests that psychedelics may exhibit state-dependent learning, and further research is required to understand how translation of learning during psychedelic states to non-psychedelic states may occur.

There are several limitations to the presented evidence. Few (n=2) human studies were identified, and none that examined changes in learning in the post-acute phase, which is of critical importance when understanding sustained behavioural changes. While one study has examined post-acute cognitive changes in humans (Wießner et al., 2022) none have been conducted for human learning. The reliance on animal models in tightly regulated contexts may not translate well into human experiences with dynamic contexts. There was considerable heterogeneity in study designs, particularly with variable dosing schedules, timings between learning and drug administrations, and training structures. Studies largely examined the acute effects of psychedelics on learning, which do not necessarily translate into a drug-free state. While there was evidence of an interaction between drug and task exposure in the post-acute phase (Šabanović et al., 2024), this requires replication. While sex differences were observed in some studies, most studies used exclusively male subjects or did not examine differences. Finally, many classical conditioning studies used repeated dosing and administration paradigms, meaning that while the first session measures acute enhancements of learning, any improvement observed on subsequent days may be contributed to by post-acute drug effects from the previous day. Additionally, such a design does not permit distinction between an acute enhancement of conditioning, and/ or an enhanced retrieval of learned associations.

In conclusion, psychedelics appear to enhance associative learning acutely, with some evidence suggesting that this may persist post-acutely. Various mechanisms were identified, including 5-HT_1A/2A_ agonism, enhanced dopamine signalling for expected rewards, and enhanced reinforcement learning rates. It seems likely that various mechanisms work synergistically to provide an enhancement for learning, underwritten by a period of enhanced neuroplasticity. Further human research is required to understand whether such learning effects are observed post-acutely. Furthermore, the directionality of the interaction between neuroplasticity and learning remains unknown. Computational modelling is emerging as a useful tool to characterise such changes and may help understand the validity of existing animal models. Further research into the context-specificity of learning changes will help shed light on how insights gained during psychotherapy may be learnt and incorporated into individuals’ lives.

**Author contributions:**

AC: conceptualisation, investigating, writing – original draft; LL: investigating, writing – reviewing and editing; FA: investigation, writing – review & editing, CB: investigation, writing – review & editing, AY: supervision, writing – review & editing, MM: conceptualisation, writing – review & editing

## Statements and declarations Ethical considerations

Ethical approval was not required.

## Consent to participate

Not applicable.

## Consent for publication

Not applicable.

## Declaration of conflicting interests

*AHY has received grant funding from LivaNova, Janssen, Compass Pathways Plc., Novartis, NIMH, CIHR, NARSAD, Stanley Medical Research Institute, MRC, Wellcome Trust. Royal College of Physicians, BMA, UBC-VGH Foundation, WEDC, CCS Depression Research Fund MSFHR, NIHR, and EU Horizon 2020, has received payments or honoraria for presentations or advisory roles from Flow Neuroscience, Novartis, Roche, Janssen, Takeda, Noema pharma, Compass Pathways Plc., AstraZeneca, Boehringer Ingelheim, Eli Lilly, LivaNova, Lundbeck, Sunovion, Servier, Janssen, Allergan, Bionomics, Sumitomo Dainippon Pharma, Sage, and Neurocentrx, and is co-editor of Journal of Psychopharmacology*.

## Funding statement

AC is a Wellcome Trust Doctoral Clinical Research Fellow (223486/Z/21/Z).

## Acknowledgements

We would like to thank Rita Moura for her thoughtful reading of the manuscript and her helpful feedback.

---

**Supplementary Table 1.**
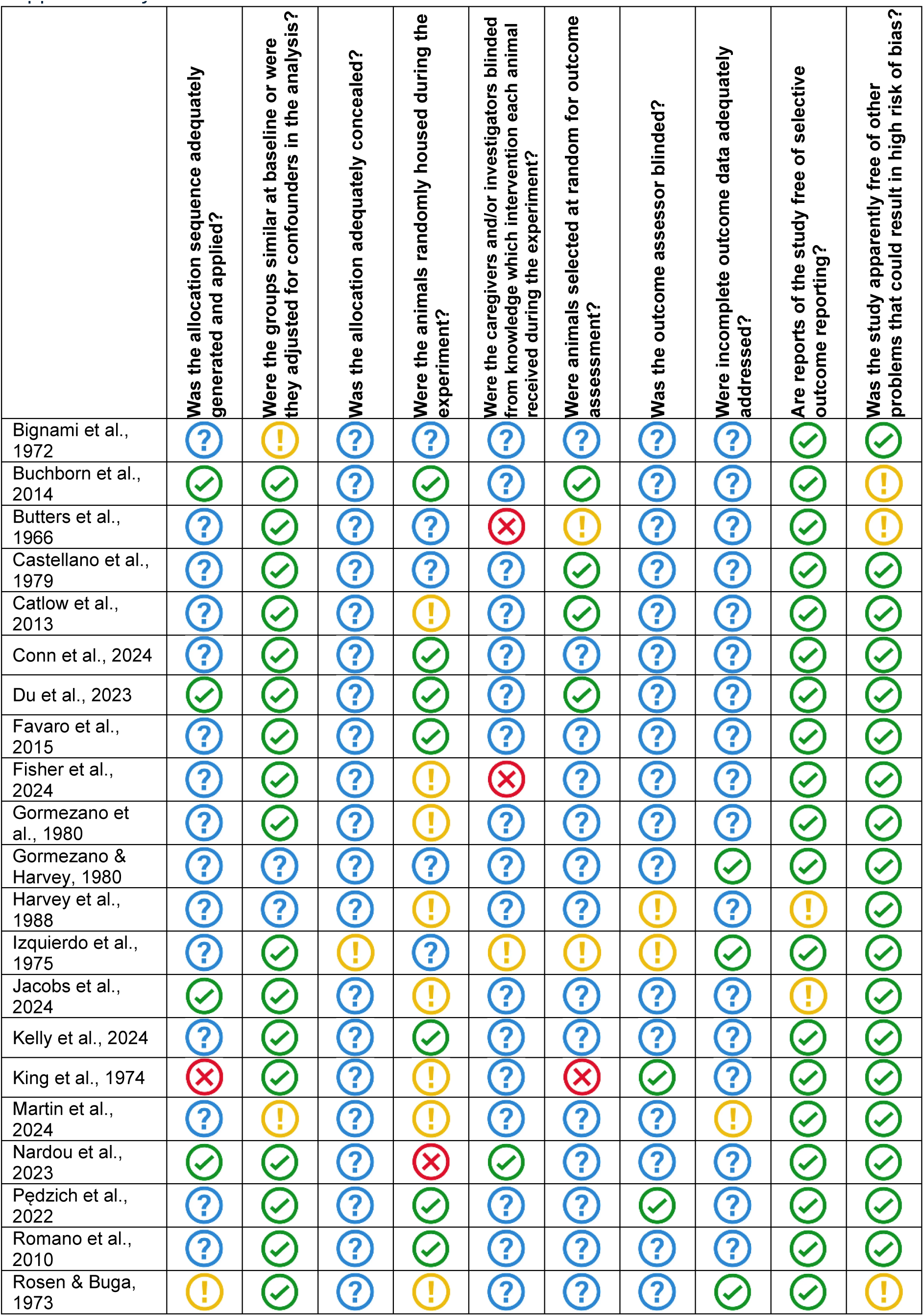
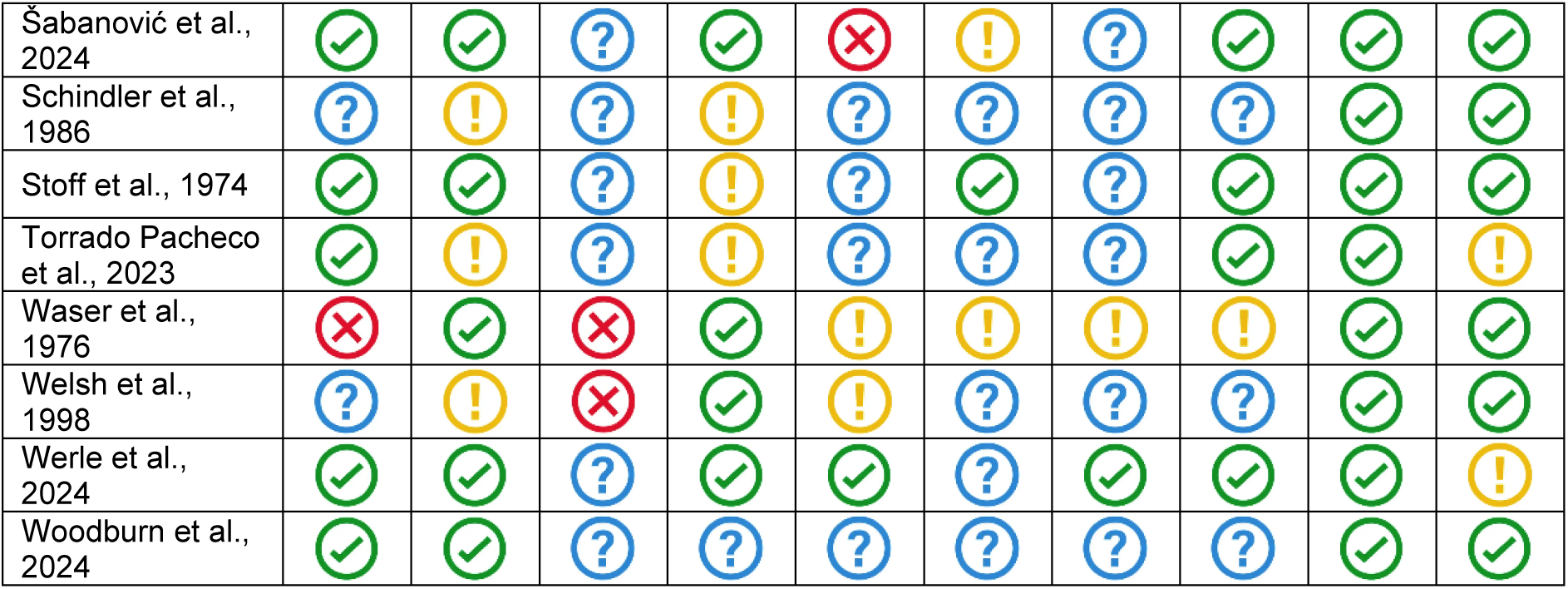
: SYRCLE’s risk of bias tool for animal studies

**Supplementary Table 2.**
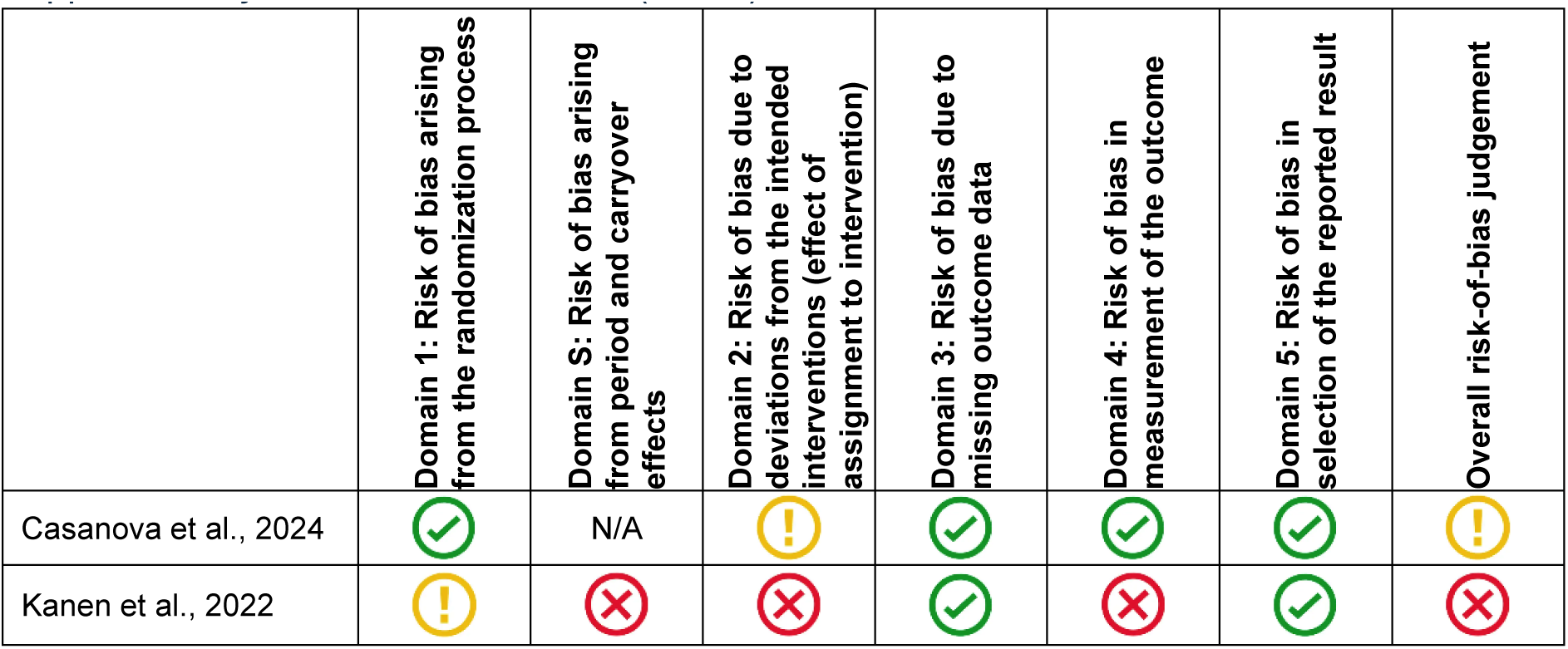
: Risk of Bias 2 (RoB 2) tool for randomised trials

## Notes

### Competing Interest Statement

MAM has research funding from Nxera and Lundbeck and received in-kind contributions from Compass Pathways. He has consulted for Boehringer Ingelheim and Nxera and received speaker fees from Takeda.
AHY has received grant funding from LivaNova, Janssen, Compass Pathways Plc., Novartis, NIMH, CIHR, NARSAD, Stanley Medical Research Institute, MRC, Wellcome Trust. Royal College of Physicians, BMA, UBC-VGH Foundation, WEDC, CCS Depression Research Fund MSFHR, NIHR, and EU Horizon 2020, has received payments or honoraria for presentations or advisory roles from Flow Neuroscience, Novartis, Roche, Janssen, Takeda, Noema pharma, Compass Pathways Plc., AstraZeneca, Boehringer Ingelheim, Eli Lilly, LivaNova, Lundbeck, Sunovion, Servier, Janssen, Allergan, Bionomics, Sumitomo Dainippon Pharma, Sage, and Neurocentrx, and is co-editor of Journal of Psychopharmacology.

